# Non-homologous DNA increases gene disruption efficiency by altering DNA repair outcomes

**DOI:** 10.1101/040212

**Authors:** CD Richardson, GJ Ray, JE Corn

**Author notes:** authors contributed equally to this work (CD Richardson, GJ Ray).

## Abstract

Cas9 endonuclease can be targeted to genomic sequences by varying the sequence of the single guide RNA (sgRNA). The activity of these Cas9-sgRNA combinations varies widely at different genomic loci and in different cell types. Thus, disrupting genes in polyploid cell lines, or using inefficient sgRNAs, can require extensive downstream screening to identify homozygous clones. We have found that linear, non-homologous oligonucleotide DNA greatly stimulates Cas9-mediated gene disruption in the absence of homology-directed repair. This stimulation greatly increases the frequency of clones with homozygous gene disruptions, even in polyploid cell lines, and rescues otherwise ineffective sgRNAs. The mechanism of enhanced gene disruption differs between human cell lines, stimulating deletion of genomic sequence and/or insertion of non-homologous oligonucleotide DNA at the edited locus in a cell line specific manner. Thus, the addition of non-homologous DNA appears to drive cells towards error-prone instead of error-free repair pathways, dramatically increasing the frequency of gene disruption.

## Main Text

Programmable genetic disruption holds great promise for the investigation of gene function and translational potential for the treatment of many diseases. Gene knockouts are commonly generated by introducing a site specific double strand break (DSB) within the gene of interest and screening for clones in which one or more alleles have been repaired in an error-prone fashion to disrupt the open reading frame^1^. The efficiency of this process is limited by the number of clones that must be screened to find the interruption, which is itself a product of the frequency of genome cutting and the frequency of disruptive repair events. The programmable Cas9 nuclease, which relies upon a targeting single guide RNA (sgRNA), has recently emerged as a popular tool for gene disruption due to its relative ease of use^2^. But Cas9-sgRNA combinations vary greatly in apparent cellular activity, from completely inactive to nearly 100% efficient, which can complicate experiments in which functional concerns place restrictions on the location to be targeted^3-6^. This variable activity has been attributed to differences in Cas9’s ability to use sgRNAs of various sequences^7,8^, but differences in the activity of a given sgRNA between cell lines and organisms suggests that location-or organism-specific modulation of DNA repair outcomes may influence observed sgRNA efficiency. Here we show that the addition of linear DNA, including non-homologous sequences, during Cas9-mediated gene ablation greatly increases the frequency of disrupting mutations in multiple human cell lines. Consequently, we dramatically increase the number of cells with homozygous gene disruptions within the edited population.

While investigating parameters to optimize rates of homology directed repair (HDR) during genome editing experiments, we found that the frequency of error-prone repair outcomes also tended to increase when single stranded HDR donor DNA was present in the editing reaction^9^. Prompted by this observation, we undertook a systematic exploration of the parameters underlying DNA-mediated stimulation of error-prone repair events. To avoid confounding effects stemming from the use of plasmid or other nucleic acid mediated delivery of Cas9, we performed editing experiments using nucleofection to directly introduce a ribonucleoprotein complex of Cas9 complexed with sgRNA (RNP) into cells^3,6^.

Targeting the EMX1 locus, we selected a sub-optimal RNP whose activity was approximately 20% in HEK293T cells., We found that the addition of a 127-mer single stranded DNA oligonucleotide derived from BFP, which lacks homology to the targeted locus and whose sequence is absent in the human genome, dramatically increased the appearance of insertions and deletions (indels) as measured by a T7E1 assay (**Figure 1A**). We henceforth refer to such non-homologous oligonucleotides as “N-oligos”. The ability of N-oligos to increase editing efficiency was titratable and depended upon oligo length, with shorter oligos losing efficacy. Native and denatured salmon sperm DNA were also capable of stimulating indels to a similar extent as synthetic single stranded oligonucleotides. Neither the charged agents Heparin and Spermidine (**Extended Data Figure 1**), nor dI-dC had little effect on editing, indicating that complex nucleic acid was necessary for stimulation. Free DNA ends were also required, as closed circular plasmid was ineffective. Importantly, DNA-mediated stimulation of indel formation is specific to the targeted site and does not increase editing at predicted off-target sites as measured by TIDE analysis^10^-^11^ (**Extended Data Figure 2**).

**• Figure 1:**
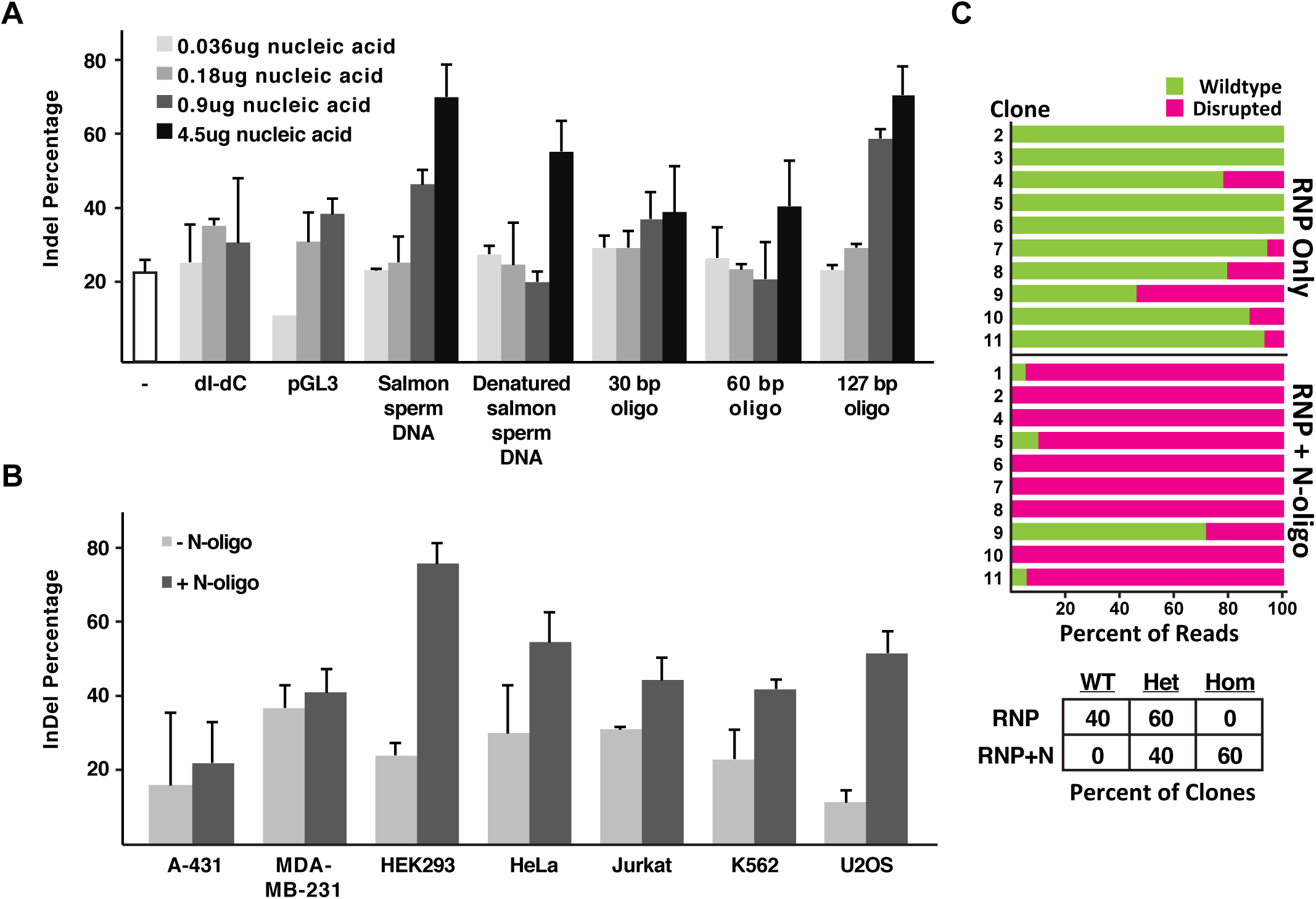
Nonhomologous DNA increases gene disruption in multiple cell types. (**A**) Single and double stranded linear nonhomologous DNA stimulates indel formation in HEK293 cells. Cas9 was targeted to the EMX1 locus with or without nucleic acid carrier agents (-, no nucleic acid). Indel formation was measured using a T7 endonuclease I assay (mean ± standard deviation of at least two independent experiments, gels presented in **Document S1**). (**B**) N-oligo DNA boosts editing rates in multiple cell types. Editing was performed as described in panel A in multiple cell types either with (dark grey bars) or without (light gray bars) 4.5ug of N-oligo. (**C**) N-oligo increases the frequency of homozygous gene disruption. HEK293 cells edited in panel B were clonally isolated and amplicons were sequenced to determine genotype. Each horizontal bar represents a single clone with green (wildtype sequence) or magenta (mutations disrupting EMX1) divisions sized according to the percentage of sequencing reads in each category. Zygosity is summarized in the lower table.

We next asked whether the use of N-oligos to increase editing could be generalized to different cell types and genomic loci. We found that N-oligos stimulated indel formation in five out of the seven cell lines tested, with tissue types ranging from bone to blood, including a five-fold increase in indels in U2OS cells **(Figure 1B)**. N-oligo stimulation of indels was observed at the YOD1 and JOSD1 loci. This stimulation at the YOD1 locus “rescued” an otherwise completely ineffective guide, more than doubling the rate of indel formation from nearly undetectable to approximately 17% (**Extended Data Figure 3**).

Because the T7E1 indel formation assay operates on an edited pool of cells and does not report on individual alleles, we used TA/TOPO cloning and Sanger sequencing to determine if increased indel frequency corresponded to a higher number of clonal homozygous knockouts. We focused on HEK293T cells, which have a tetraploid genome and are thus a stringent test case for the formation of homozygous knockouts. Characterizing clonally isolated edited cells, we found that editing with RNP alone yielded 40% heterozygous clones and no homozygous knockouts, whereas RNP with N-oligos yielded 40% heterozygotes and 60% homozygous knockouts (**Figure 1C**). Hence, the use of N-oligos is a simple and effective technique to increase the frequency of homozygous gene disruption.

Sequence analysis of the alleles in HEK293 editing reactions revealed that N-oligo treatment increased the rate of both insertions and deletions relative to RNP treatment alone (**Figure 2A**). These indels included simple deletion of sequence around the cut site and insertion of random sequence, but surprisingly also insertion of N-oligo sequence and insertion of the DNA template used for *in vitro* transcription of the sgRNA (example indels, **Figure 2B**; all indels **Extended Data Figure 4**). The frequent presence of sgRNA template sequence was particularly striking, as the N-oligo was approximately 1000-fold more abundant in the nucleofection reactions (**Extended Data Figure 5**). The occasional insertion of non homologous DNA into double strand breaks has been reported in yeast^12-14^ and mice^15-17^ and the insertion of short phosphorothioate-protected oligos forms the basis of the GUIDE-Seq method to detect off-target genome editing events^18^, but the ability of non-homologous single stranded DNA to greatly increase gene knockout by stimulating these events is surprising and to the best of our knowledge unprecedented.

**• Figure 2:**
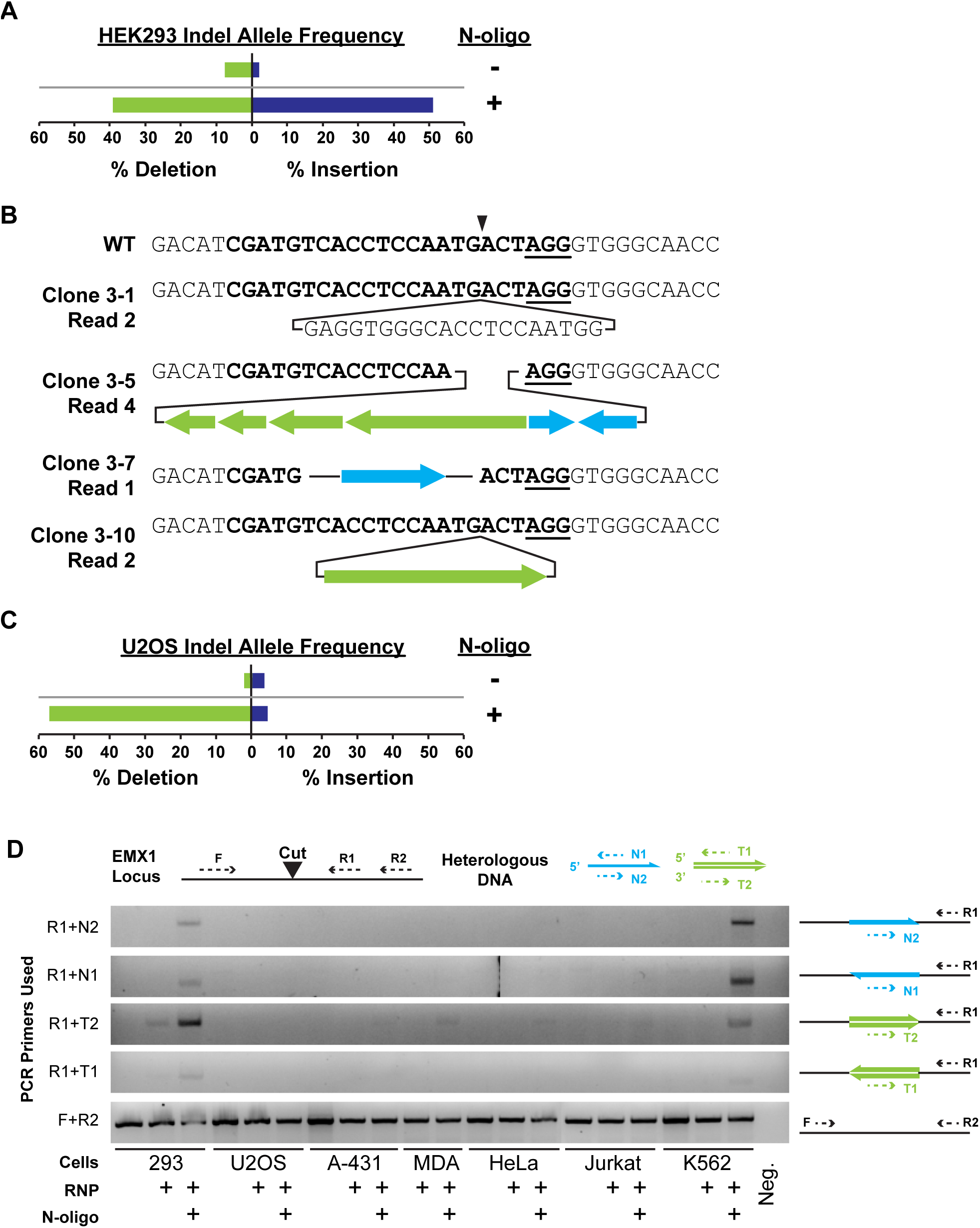
N-oligo stimulation of gene disruption promotes error-prone repair events. (**A**) N-oligo stimulates insertions and deletions in HEK293 cells. The allele frequency of deletions (left, green) and insertions (blue, right) are shown for nucleofections performed with or without N-oligo. Editing is summarized for each clone in **Extended Data Figure 4**. Raw data is available in **Document S3**. (**B**) Inserted sequences are derived from single and double stranded heterologous DNA. Wildtype EMX1 sequence is presented at top with the protospacer (bold), PAM (bold underline), and cut site (triangle) diagrammed. Four example alleles are presented below with sgRNA template (green) and/or N-oligo (blue) sequence inserted in both orientations. Complete sequencing alignments are available in Document S3. (**C**) N-oligo stimulates deletions in U2OS cells. Multiple sequence reads from edited cell populations are presented as described in **Figure 2A**. (**D**) Insertion of nonhomologous DNA primarily occurs in HEK293 and K562 cells. DNA harvested from edited cell populations was evaluated using a panel of PCR reactions (diagrammed at top). Primers N1 and N2 anneal to the N-oligo sequence; primers T1 and T2 anneal to residual sgRNA template.

Our observations of sequence insertion in HEK293T cells motivated us to investigate the nature of N-oligo stimulated indel formation in other cell types. Surprisingly, Sanger sequencing of U2OS editing outcomes revealed that N-oligo treatment primarily stimulated the appearance of large deletions, and not insertions, as compared to RNP alone editing (**Figure 2C**). To determine the propensity of various cell types to insert sequences into a Cas9 break point, we designed a PCR assay to amplify either the N-oligo or sgRNA transcription template in various orientations. This assay confirmed sequence insertion in K562 and HEK293T cells, but not in any of the other cell lines that show robust N-oligo indel stimulation (**Figures 2D, 1B**). The ability of N-oligos to stimulate gene disruption with such different molecular outcomes suggests that they stimulate classical or alternative end-joining pathways^19^, thereby boosting the rate of error-prone DSB repair.

Given the large excess of N-oligo over sgRNA template, we wondered if providing high concentrations of double stranded non-homologous DNA would also effectively stimulate insertion in these cells. We tested both single and double stranded N-oligos for their potential to increase indels at the EMX1 locus, double purifying the sgRNA before use to ensure that the double stranded sgRNA template was completely removed. We found that double stranded N-oligo stimulated indels, though about twofold less effectively than single stranded N-oligo (**Extended Data Figure 6A**). Using a PCR assay for sequence integration, we found no evidence of sgRNA template insertion in doubly-purified samples, but greater integration of the duplex N-oligo relative to single-stranded N-oligo (**Extended Data Figure 6B**). This result suggests that explicitly designing a duplex N-oligo could further bias cells towards a specific outcome, for example inserting a cassette that encodes stop codons in multiple frames and orientations may further bias cells towards a specific repair outcome. This strategy has been proposed for HDR-mediated gene disruption, but to out knowledge has not been attempted with non-homologous integration of oligonucleotides^20^.

Taken together, our data support a model in which cells fidelitously repair most Cas9-generated DSBs using error-free repair pathways that do not produce measurable indels, but occasional error-prone repair causes indels that ablate portions of the Cas9 protospacer and/or PAM, thereby preventing further cutting and producing a measurable outcome (**Extended Data Figure 7**). We note that if repair outcomes depend upon sequence context, this activity could cause Cas9-sgRNA combinations with high *in vitro* activity to display poor cellular activity or for sgRNA activity to differ between cell lines^7,8^. The addition of N-oligo during editing appears to stimulate error-prone end-joining pathways that differ among cell types (e.g. end-joining of exogenous nucleic acid in HEK293T and large deletions in U2OS) but have the net effect of increasing the rate of gene disruption. We anticipate that the use of N-oligos will be extremely valuable in generating homozygously gene-disrupted cell lines or organisms, and will be particularly effective in challenging polyploid contexts.

## Supplemental Documents

- **Document S1:** Uncropped gels from Figures 1A, 1B, 2D, and Extended 6.
- **Document S2:** Sequences of constructs/PCR primers/etc
- **Document S3:** Sequences for all clones/reads

## Materials and Methods

### Cell Lines and Cell Culture

A-431, HEK293, HeLa, Jurkat, K562, MDA-MB-231, and U2OS cells were acquired from the UC Berkeley Tissue Culture Facility. A-431, HeLa, and MDA-MB-231 cells were maintained in DMEM glutamax medium supplemented with 10% fetal bovine serum, 1% sodium pyruvate, 1% non-essential amino acids, and 100 ug/mL penicillin-streptomycin. HEK293 and U2OS cells were maintained in DMEM medium supplemented with 10% fetal bovine serum, 1% sodium pyruvate, and 100 ug/mL penicillin-streptomycin. Jurkat and K562 cells were maintained in RPMI medium supplemented with 10% fetal bovine serum, 1% sodium pyruvate, and 100 ug/mL penicillin-streptomycin.

### Cas9 and RNA Preparation

*S. pyogenes* Cas9 (pMJ915, Addgene #) with two nuclear localization signal peptides and an HA tag at the C-terminus were purified by a combination of affinity, ion exchange, and size exclusion chromatography steps as described^21^, except protein was eluted at 40uM in 20 mM HEPES KOH pH 7.5, 5% glycerol, 150 mM KCl, 1 mM DTT.

sgRNAs were generated by HiScribe (NEB E2050S) T7 in vitro transcription using PCR-generated DNA as a template (dx.doi.org/10.17504/protocols.io.dm749m). Complete sequences for all sgRNA templates can be found in Document S2.

### Cas9 RNP Assembly and Nucleofection

100 pmoles of Cas9-2NLS was diluted to a final volume of 5uL with Cas9 buffer (20 mM HEPES (pH 7.5), 150 mM KCl, 1 mM MgCl2, 10% glycerol and 1 mM TCEP) and mixed slowly into 5uL of Cas9 buffer containing 120 pmoles of L2 sgRNA. The resulting mixture was incubated for ten minutes at room temperature to allow RNP formation. 2E+05 cells were harvested, washed once in PBS, and resuspended in 20uL of nucleofection buffer (Lonza, Basel, Switzerland). 10uL of RNP mixture, 4.5 uL of N-oligo, and cell suspension were combined in a Lonza 4d strip nucleocuvette. Reaction mixtures were electroporated, incubated in the nucleocuvette at RT for ten minutes, and transferred to culture dishes containing pre-warmed media (dx.doi.org/10.17504/protocols.io.dm649d). Editing outcomes were measured two days post-nucleofection by T7E1 (see below). Resuspension buffer and electroporation contions were the following for each cell line: A-431 in SF with EQ-100, HEK293 in SF with DS-150, HeLa in SE with CN-114, Jurkat in SE with CL-120, K562 in SF with FF-120, MDA-MB-231 in SE with CH-125, and U2OS in SE with CM104.

### PCR Amplification of Edited Regions

PCR amplification of EMX1 was done using primers oCR295 and oCR296. PCR amplification of YOD1 was done using YOD1f and YOD1r. PCR amplification of JOSD1 was done using JOSD1f and JOSD1r. PCR reactions were performed with 200 ng of genomic DNA and Kapa Hot Start high-fidelity polymerase with the GC buffer. The thermocycler was set for one cycle of 95°C for 5 min, 30 cycles of 98°C for 20 sec, 62°C for 15 sec, 72°C for 30 sec, and one cycle of 72°C for 1 min, and held at 4°C.

### T7EI Assay

The rate of Cas9 mediated gene disruption was measured by T7 endonulcease I digestion of hybridized PCR. 200 ng of PCR DNA in 1X NEB Buffer 2 was hybridized in a thermocycler under the following conditions: 95°C for 5 min, 95-85°C at -−2°C/sec, 85-25°C at -.1°C/sec, and held at 4°C. 10 units of T7EI (NEB, M0302) were added to the sample and was incubated at 37°C for 15 min. The sample was then immediately run on a 2% agarose gel containing ethidium bromide. Band intensities were quantified by imageJ. Indel percentage was calculated using the following equation: (1-(1-(cut product intensities/ uncut + cut product intensities))1/2) x100^5^.

### Insert Based PCR Assay

To assay the insertion of N-oligo and sgRNA template DNA into the cut site, a reverse primer (oGJR102) was designed to pair with forward primers homologous to the: BFP N-oligo inserted in the forward direction (oGJR097), BFP N-oligo inserted in the reverse direction (oGJR098), the T7 promoter of the sgRNA template inserted in the forward direction (oGJR099), and the T7 promoter of the sgRNA template inserted in the reverse direction (oGJR100). Presence of EMX1 DNA in the PCR reaction was verified using the EMX1 PCR performed above. PCR reactions were performed with 200 ng of genomic DNA and Kapa Hot Start high-fidelity polymerase. The thermocycler was set for one cycle of 95°C for 5 min, 30 cycles of 98°C for 20 sec, 64°C for 15 sec, 72°C for 30 sec, and one cycle of 72°C for 1 min, and held at 4°C. The sample was then run on a 2% agarose gel containing ethidium bromide.

### qPCR on RNP

sgRNA template and N-oligo DNA present in the nucleofection reaction mixture was quantified on a Mastercycler 2 qPCR machine (Eppendorf, Hamburg). Primers oCR427 and oCR428, sgRNA template; oGJR103 and oGJR104, N-oligo; **Document S2** were used at a final concentration of 500nM in Power SYBR green reaction mixture (Thermo Fisher). Reaction conditions were 95°C for 10 minutes followed by 40 cycles of 95°C for 30 seconds and 65°C for 60 seconds. The ratio of N-oligo to sgRNA template was quantified using the equation r=2^(Ct_sgRNA_-Ct_N-oligo_). Ratios from three serial dilutions of template DNA were averaged and presented as mean ± standard deviation.

### TIDE Analysis

Off-target analysis was performed on the top 4 targets given by the online tool available at http://crispr.mit.edu/22. Sequences are presented in **Extended Data Figure 2** and **Document S2**. Genomic sequences were amplified using KAPA polymerase and Sanger sequenced by the UC Berkeley Sequencing Core. PCR reactions were performed using 200 ng of genomic DNA and the thermocycler was set for one cycle of 95°C for 5 min, 30 cycles of 98°C for 20 sec, the annealing temperature for 15 sec, 72°C for 30 sec, and one cycle of 72°C for 1 min, and held at 4°C. Off-Target 1 PCR was done with oGJR051 and oGJR096 with a 64°C annealing temperature, off-Target 2 PCR was done with oGJR090 and oGJR091 with a 62°C annealing temperature, off-Target 3 PCR was done with oGJR053 and oGJR071 with a 64°C annealing temperature, and off-Target 4 PCR was done with oGJR054 and oGJR072 with a 66°C annealing temperature. ABIF files were uploaded to http://www.tide.nki.nl for analysis^11^.

## Acknowledgments

We thank members of the Corn lab for helpful discussions on this manuscript. This work was supported by the Li Ka Shing Foundation.

---

**Extended Data Figure 1:**
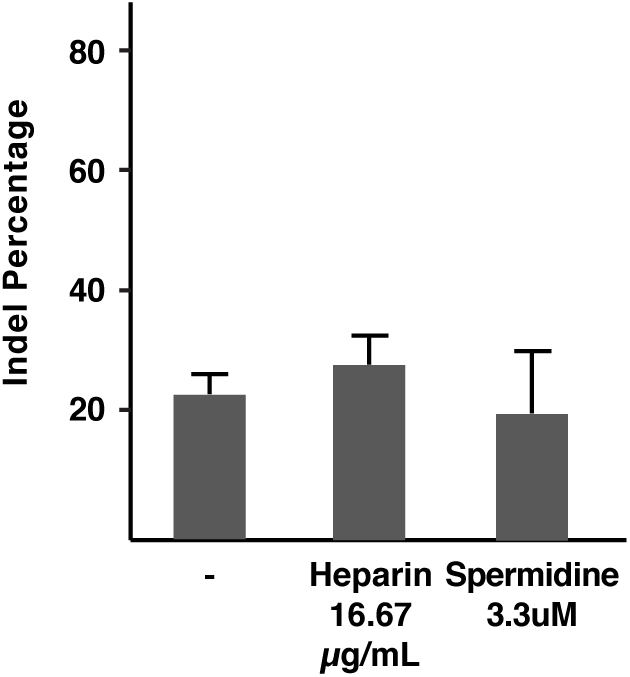
Heparin and spermidine do not stimulate editing at the EMX1 locus in HEK293T cells. Editing was performed as described in Figure 1A.

**Extended Data Figure 2:**
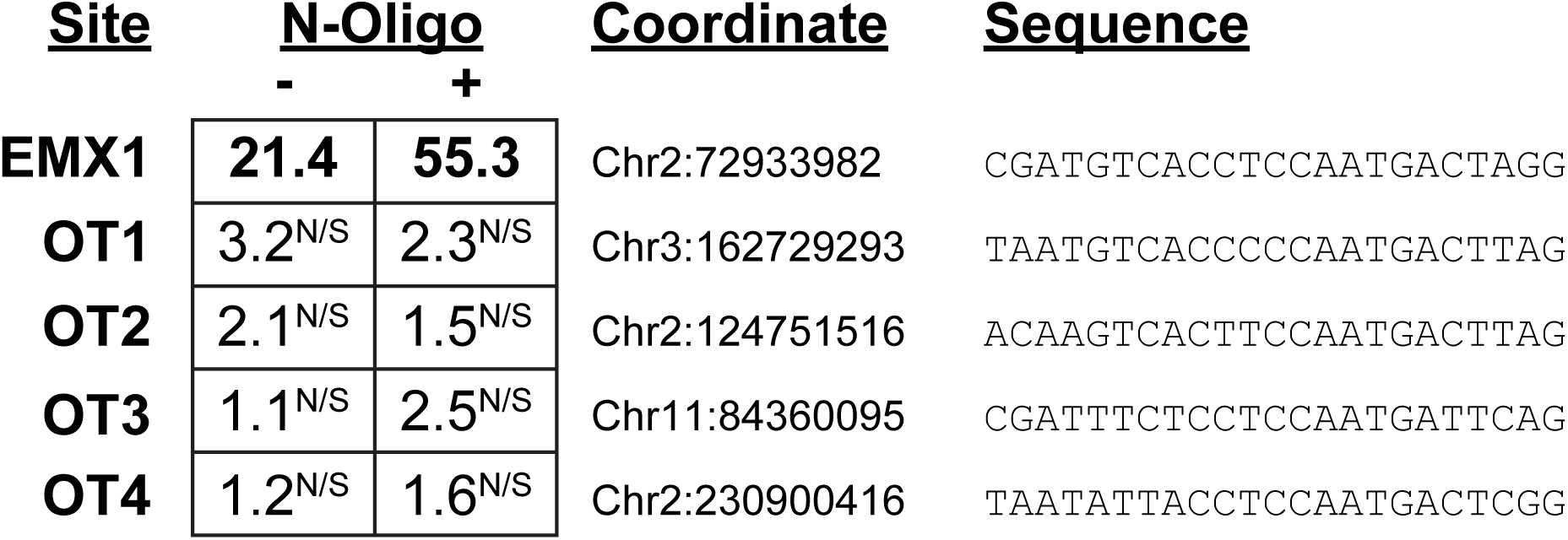
N-oligo does not stimulate measurable editing at predicted off-target sites. TIDE analysis^10^ was performed at EMX1 and four predicted off-target sites^20^ following editing experiments performed in the presence or absence of 4.5ug of salmon sperm DNA. Total efficiency is presented for each case. N/S – not significant, apparent editing at this locus was not significant relative to background.

**Extended Data Figure 3:**
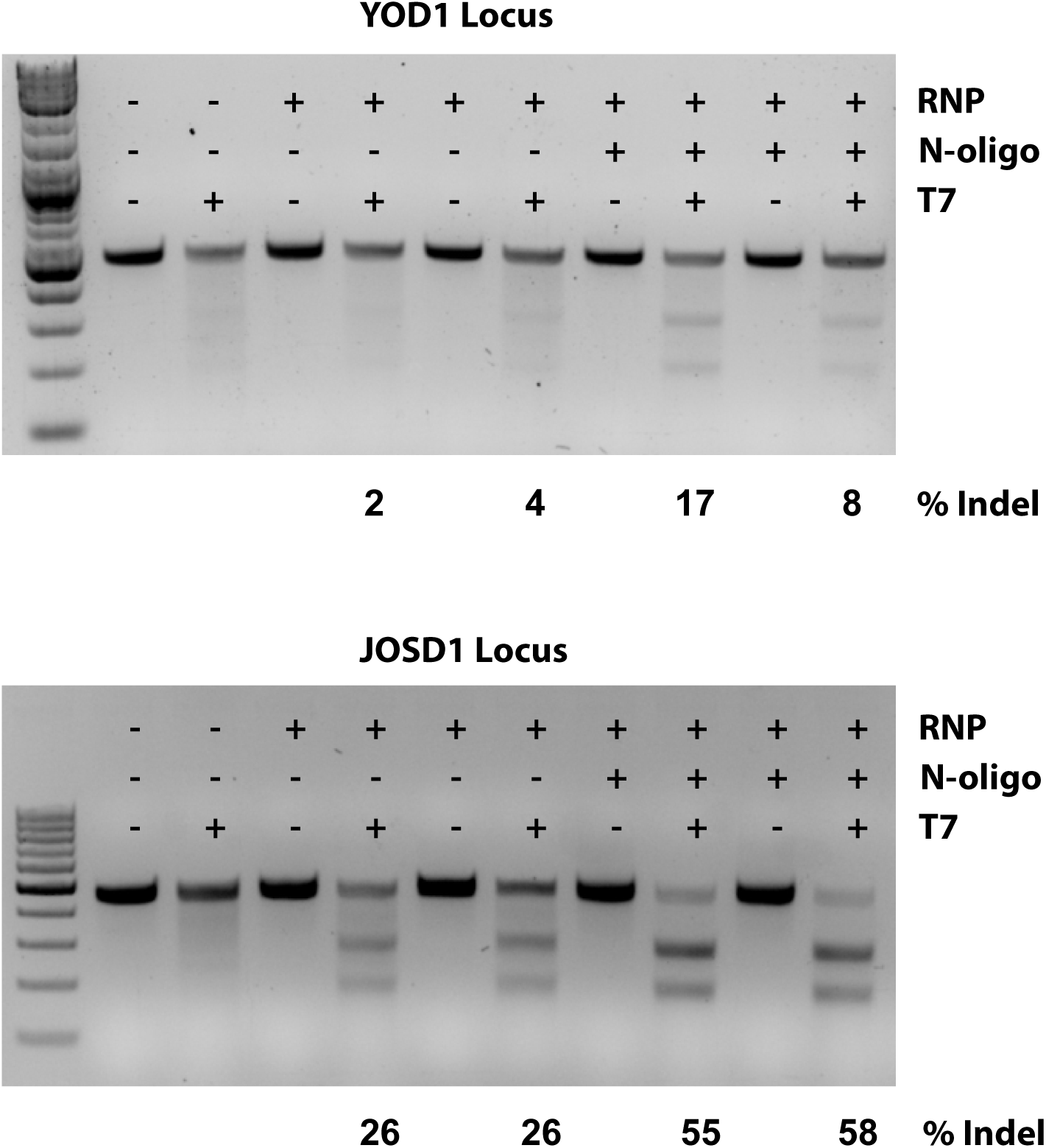
N-oligo stimulates editing at the YOD1 and JOSD1 loci in HEK293 cells. Editing experiments were performed with or without N-oligo as indicated. Indel percentage was assayed by T7 endonuclease cleavage and gel densitometry.

**Extended Data Figure 4:**
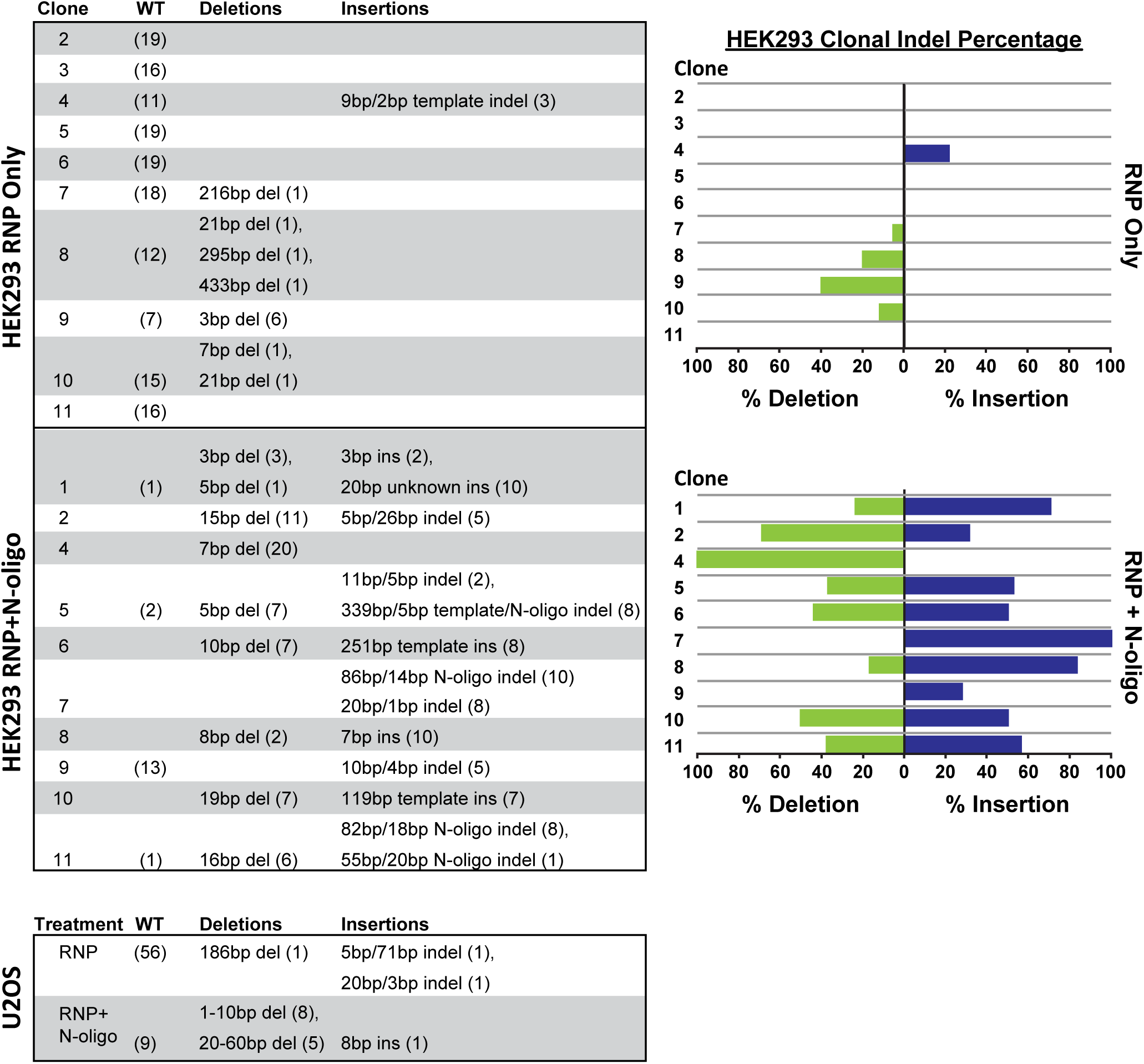
N-oligo stimulates insertions and deletions in HEK293 cells. Sequence reads from clonal cell populations were binned into three categories (WT, unmodified; deletions, clear removal of sequence; and insertions, added sequence with or without flanking deletions) and presented in table form. The complete sequence of each read can be found in Document S3. Bar graphs present clonal indel percentages.

**Extended Data Figure 5:**
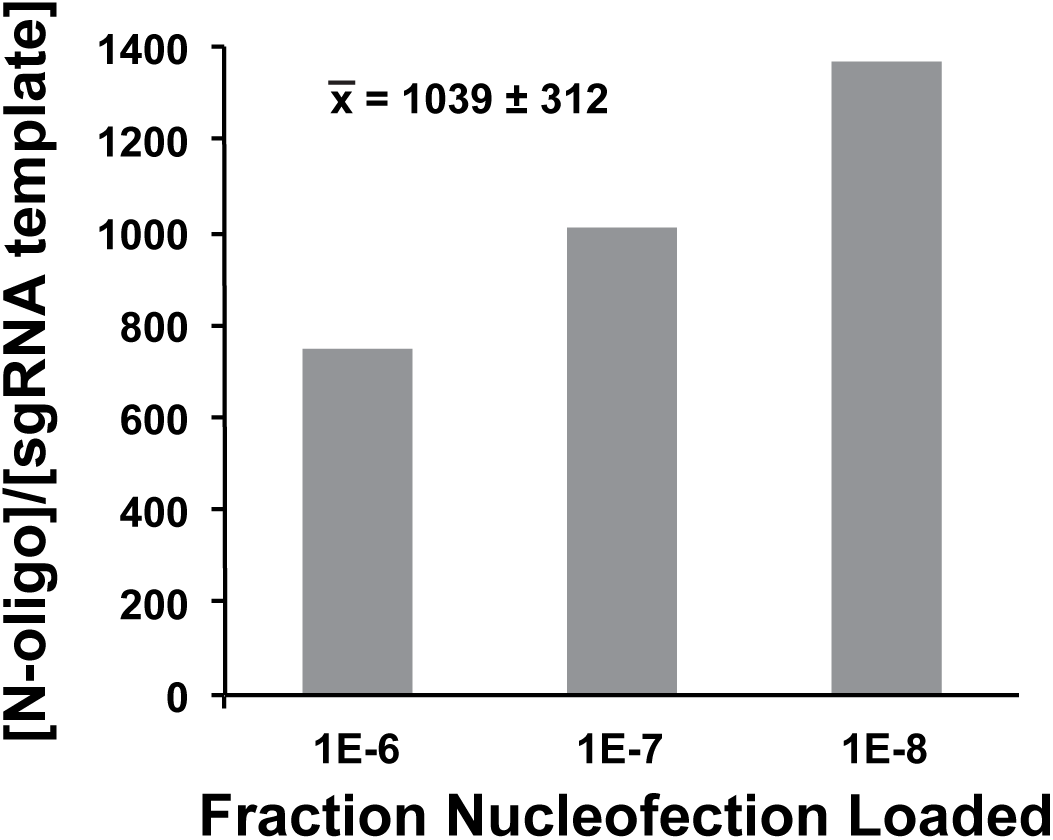
N-oligo is present in excess of sgRNA template. Nucleofection mixtures were serially diluted and the abundance of N-oligo or sgRNA template were quantified by qPCR. Fold enrichment of N-oligo over sgRNA template are presented for three serial dilutions of nucleofection mixtures. Inset number is the mean +/- standard deviation of these three values.

**Extended Data Figure 6:**
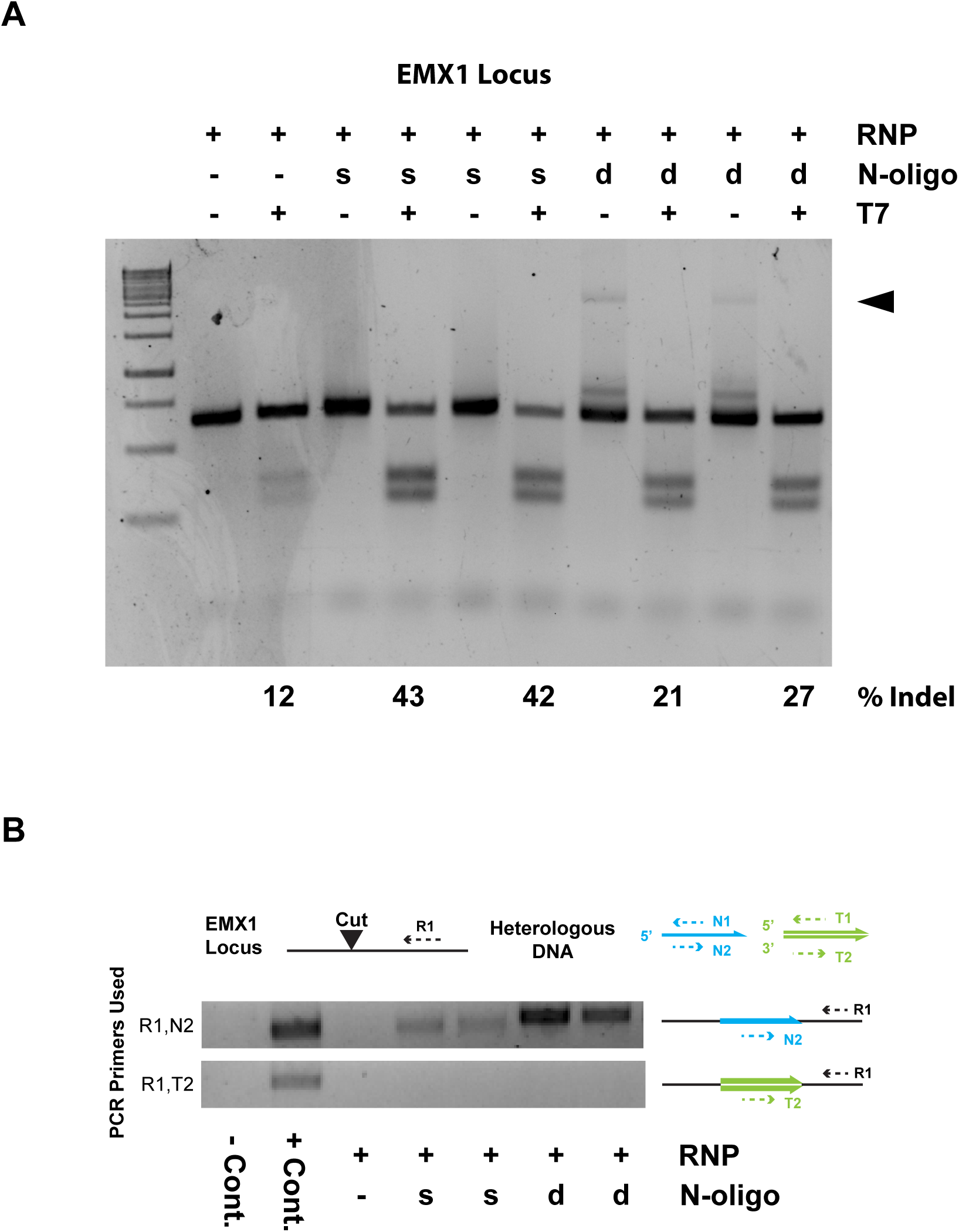
sgRNA template DNA is not required for N-oligo increases to gene disruption. (A) Extensively purified sgRNA (free of sgRNA template DNA) was used in editing experiments at the EMX1 locus. T7 editing rates increased dramatically with the addition of single or double stranded DNA. (B) PCR for N-oligo or sgRNA template insertion indicates that insertion events are derived from the N-oligo rather than the sgRNA template. Double stranded N-oligos (d) tend to insert more efficiently than single stranded N-oligos (s).

**Extended Data Figure 7:**
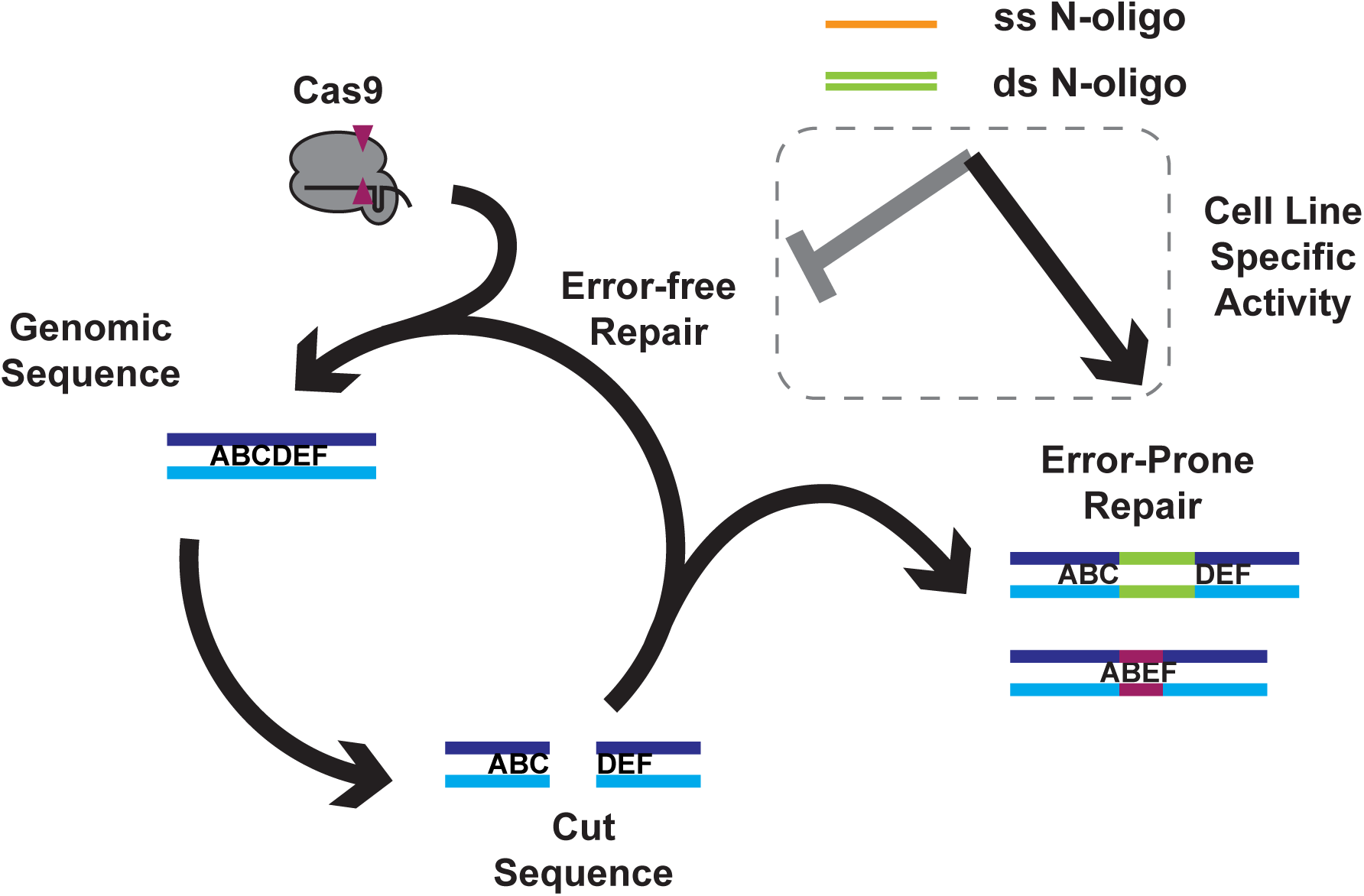
Model for N-oligo action. Cas9-sgRNA (grey) recognizes and cuts genomic sequence (ABCDEF). Cellular repair processes reseal most breaks in an error-free fashion, which restores the Cas9 recognition sequence and permits additional rounds of cutting. Error-prone repair events such as insertions (green sequence) and deletions (maroon region) disrupt the Cas9 recognition sequence and prevent cutting. Single and double stranded N-oligo act to inhibit error-free repair and promote error-prone repair events. Error-prone repair events stimulated by N-oligo are cell-line dependent.

## References

1. Yang, L., Yang, J. L., Byrne, S., Pan, J. & Church, G. M. CRISPR/Cas9-Directed Genome Editing of Cultured Cells. Current protocols in molecular biology / edited by Frederick M Ausubel [et al] 107, 31.1.1–31.1.17 (2014).

2. Doudna, J. A. & Charpentier, E. Genome editing. The new frontier of genome engineering with CRISPR-Cas9. Science 346, 1258096 (2014).

3. Lin, S., Staahl, B. T., Alla, R. K. & Doudna, J. A. Enhanced homology-directed human genome engineering by controlled timing of CRISPR/Cas9 delivery. eLife 4, (2014).

4. Yang, L. et al. Optimization of scarless human stem cell genome editing. Nucleic Acids Research 41, 9049–9061 (2013).

5. Ran, F. A. et al. Double nicking by RNA-guided CRISPR Cas9 for enhanced genome editing specificity. Cell 154, 1380–1389 (2013).

6. Kim, S., Kim, D., Cho, S. W., Kim, J. & Kim, J.-S. Highly efficient RNA-guided genome editing in human cells via delivery of purified Cas9 ribonucleoproteins. Genome Research 24, 1012–1019 (2014).

7. Doench, J. G. et al. Rational design of highly active sgRNAs for CRISPR-Cas9-mediated gene inactivation. Nat Biotechnol 32, 1262–1267 (2014).

8. Moreno-Mateos, M. A. et al. CRISPRscan: designing highly efficient sgRNAs for CRISPR-Cas9 targeting in vivo. Nat Meth 12, 982–988 (2015).

9. Richardson, C. D., Ray, G. J., DeWitt, M. A., Curie, G. L. & Corn, J. E. Enhancing homology-directed genome editing by catalytically active and inactive CRISPR-Cas9 using asymmetric donor DNA. Nat Biotechnol (2016). doi:10.1038/nbt.3481

10. Hsu, P. D. et al. DNA targeting specificity of RNA-guided Cas9 nucleases. Nat Biotechnol 31, 827–832 (2013).

11. Brinkman, E. K., Chen, T., Amendola, M. & van Steensel, B. Easy quantitative assessment of genome editing by sequence trace decomposition. Nucleic Acids Research 42, e168 (2014).

12. Moore, J. K. & Haber, J. E. Capture of retrotransposon DNA at the sites of chromosomal double-strand breaks. Nature 383, 644–646 (1996).

13. Teng, S. C., Kim, B. & Gabriel, A. Retrotransposon reverse-transcriptase-mediated repair of chromosomal breaks. Nature 383, 641–644 (1996).

14. Haviv-Chesner, A., Kobayashi, Y., Gabriel, A. & Kupiec, M. Capture of linear fragments at a double-strand break in yeast. Nucleic Acids Research 35, 5192–5202 (2007).

15. Lin, Y. & Waldman, A. S. Promiscuous patching of broken chromosomes in mammalian cells with extrachromosomal DNA. Nucleic Acids Research 29, 3975–3981 (2001).

16. Onozawa, M. et al. Repair of DNA double-strand breaks by templated nucleotide sequence insertions derived from distant regions of the genome. Proceedings of the National Academy of Sciences 111, 7729–7734 (2014).

17. Ono, R. et al. Double strand break repair by capture of retrotransposon sequences and reverse-transcribed spliced mRNA sequences in mouse zygotes. Sci Rep 5, 12281 (2015).

18. Tsai, S. Q. et al. GUIDE-seq enables genome-wide profiling of off-target cleavage by CRISPR-Cas nucleases. Nat Biotechnol 33, 187–197 (2015).

19. Sfeir, A. & Symington, L. S. Microhomology-Mediated End Joining: A Back-up Survival Mechanism or Dedicated Pathway? Trends Biochem Sci 40, 701–714 (2015).

20. Gagnon, J. A. et al. Efficient mutagenesis by Cas9 protein-mediated oligonucleotide insertion and large-scale assessment of single-guide RNAs. PLoS ONE 9, e98186 (2014).

21. Jinek, M. et al. A Programmable Dual-RNA-Guided DNA Endonuclease in Adaptive Bacterial Immunity. Science 337, 816–821 (2012).

22. Hsu, P. D., Lander, E. S. & Zhang, F. Development and applications of CRISPR-Cas9 for genome engineering. Cell 157, 1262–1278 (2014).

